# PAN-GWES: Pangenome-spanning epistasis and co-selection analysis via de Bruijn graphs

**DOI:** 10.1101/2023.09.07.556769

**Authors:** Juri Kuronen, Samuel Horsfield, Anna K. Pöntinen, Sergio Arredondo-Alonso, Harry Thorpe, Rebecca Gladstone, Rob J.L. Willems, Stephen D. Bentley, Nicholas J Croucher, Johan Pensar, John A. Lees, Gerry Tonkin-Hill, Jukka Corander

**Affiliations:** Department of Biostatistics, University of Oslo, Oslo, Norway; MRC Centre for Global Infectious Disease Analysis, Department of Infectious Disease Epidemiology, Imperial College London, London, United Kingdom; European Bioinformatics Institute, Cambridge, United Kingdom; Norwegian National Advisory Unit on Detection of Antimicrobial Resistance, Department of Microbiology and Infection Control, University Hospital of North Norway, Tromsø, Norway; Parasites and Microbes, Wellcome Sanger Institute, Cambridge, United Kingdom; Department of Medical Microbiology, University Medical Center Utrecht, Utrecht, Netherlands; Department of Mathematics, University of Oslo, Oslo, Norway; Helsinki Institute of Information Technology, Department of Mathematics and Statistics, University of Helsinki, Helsinki, Finland

**Author notes:** These authors contributed equally (John A. Lees, Gerry Tonkin-Hill, Jukka Corander).

**Keywords:** pangenome, microbial genomics, GWAS, epistasis

## Abstract

Studies of bacterial adaptation and evolution are hampered by the difficulty of measuring traits such as virulence, drug resistance and transmissibility in large populations. In contrast, it is now feasible to obtain high-quality complete assemblies of many bacterial genomes thanks to scalable and affordable long-read sequencing technology. To exploit this opportunity we introduce a phenotype- and alignment-free method for discovering co-selected and epistatically interacting genomic variation from genome assemblies covering both core and accessory parts of genomes. Our approach uses a compact coloured de Bruijn graph to approximate the intra-genome distances between pairs of loci for a collection of bacterial genomes to account for the impacts of linkage disequilibrium (LD). We demonstrate the versatility of our approach to efficiently identify associations between loci linked with drug resistance and adaptation to the hospital niche in the major human bacterial pathogens *Streptococcus pneumoniae* and *Enterococcus faecalis*.

## Introduction

Epistatic interactions between polymorphisms in DNA are recognized as important drivers of evolution in numerous organisms. It was recently established that even weak to moderate co-selective and epistatic effects can firmly manifest themselves in bacteria, in contrast with eukaryotes, where only stronger effects bring sufficient selective advantages (1). The rapidly increasing amount of whole-genome data for many named species of bacteria has opened up a possibility to identify such effects on a pan-genomic level. Recent approaches to genome-scale analysis of co-variation at single-nucleotide resolution, termed as genome-wide epistasis and co-selection study (GWES), have demonstrated ample potential to uncover drivers of adaptation, virulence, survival and antimicrobial resistance from densely sampled populations of major pathogens (2–5).

The GWES approach can be considered as a phenotype-free biological hypothesis generator that works in a complementary manner compared with genome-wide association study (GWAS), which also aims to generate hypotheses of causal drivers. GWAS works by correlating genomic variation with measured phenotypic variation, and has been widely applied to the study of bacterial traits, most notably antibiotic resistance (6–8). The availability of standardised and accurate bacterial phenotyping is often limited, focusing primarily on specific phenotypes like antibiotic resistance, as more opaque phenotypes like transmissibility, hospital adaptation, and virulence are harder to quantify. GWES holds considerable promise to address these difficulties, by directly measuring co-evolutionary signals, driving molecular discoveries. Existing GWES approaches have predominantly relied upon multiple sequence alignments produced by mapping reads to a single reference genome. This neglects signals found in accessory elements that are absent from the reference genome, including short indels and variation found in accessory genes. Additionally, missing data in alignments can be difficult to handle correctly, and genotyping quality can vary between samples due to the choice of thresholds used for filtering. To overcome these shortcomings and expand the potential of GWES, we developed an alignment-free approach using coloured de Bruijn graphs to distinguish between short- and long-distance linkage disequilibrium in bacterial genomes.

Use of de Bruijn graphs in genome assembly, reference genome representation and variant calling has become commonplace in population genomics, in particular for bacteria (9–11). A coloured and compacted de Bruijn graph offers a powerful representation of variation present across a collection of genome assemblies that can be utilised for a wide range of applications, including population structure analysis (12) and GWAS (7, 13) and genotyping (14). Within a coloured compacted de Bruijn graph, variable length sequences or ‘unitigs’ represent loci within each genome, with the colour indicating which genome the unitig belongs to. As a measure of genomic variation within a population, unitigs are advantageous as they can efficiently tag a range of different types of changes, including mutations, indels and variation in gene content, all of which represent important processes in evolution and adaptation of bacteria. Nevertheless, obtaining a coloured and compacted de Bruijn graph for a large set of input genome assemblies is an intensive computational problem, which has only recently become accessible due to algorithmic advances (9, 15, 16). One recent coloured de Bruijn graph construction algorithm, Cuttlefish, provides an optimal starting point for a pangenomic GWES that does not rely on a single reference genome based representation of variation (15). Using this data structure, we have developed an efficient algorithm, PAN-GWES, that computes approximate distances between loci within a pangenome and estimates the signal of epistasis or co-selection between loci using Mutual Information (MI). Applications to large pneumococcal and *E. faecalis* genomic datasets, revealed not only previously identified but also new indications of co-selection between genomic loci, linked to antibiotic resistance and adaptation to hospital environments.

## Results

### Overview

Our PAN-GWES algorithm starts by constructing a coloured, compacted de Bruijn graph for each genome assembly, as depicted in Figure 1. Figure 2 provides a comprehensive overview of the various stages and adjustable parameters within the algorithm.

**Fig. 1.**
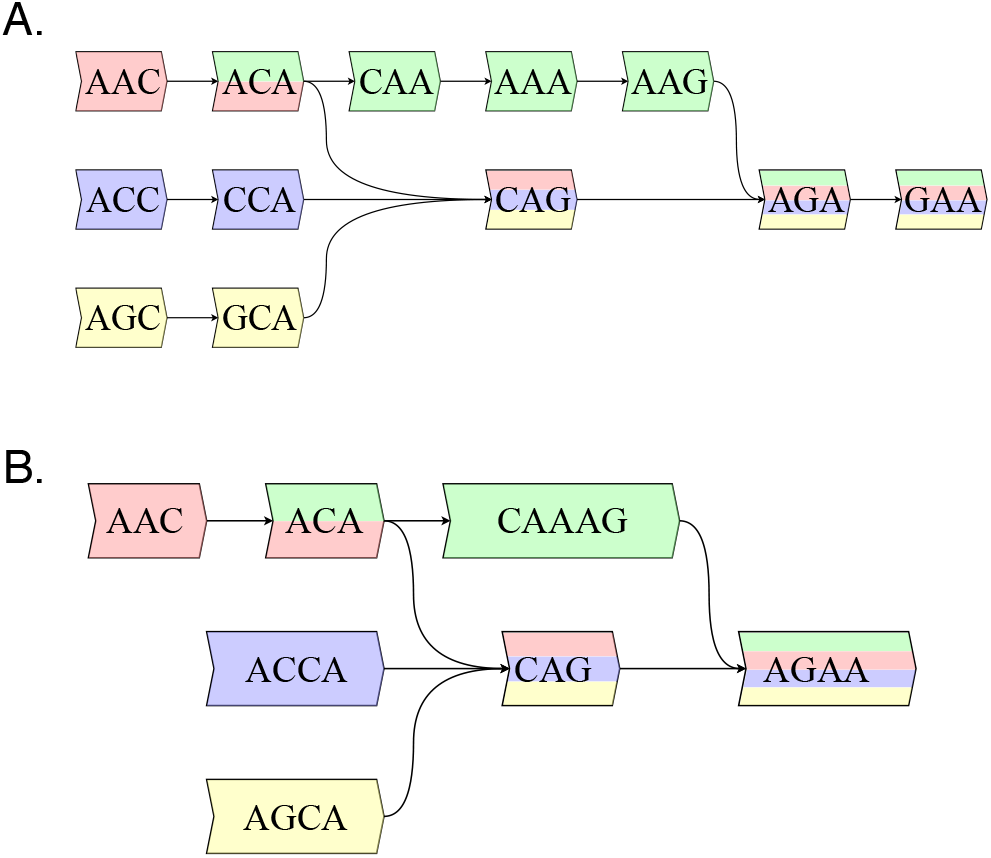
A) An uncompacted de Bruijn graph for four example sequences, S=AGCAGAA, ACCAGAA, ACAAAGAA, AACAGAA with a kmer length of 3. The hexagons are the vertices, which are the canonical 3-mers contained in S, and two vertices v and w connected by an edge if there exists a 4-mer in S whose prefix is v and suffix w. B) The corresponding compacted de Bruijn graph.

**Fig. 2.**
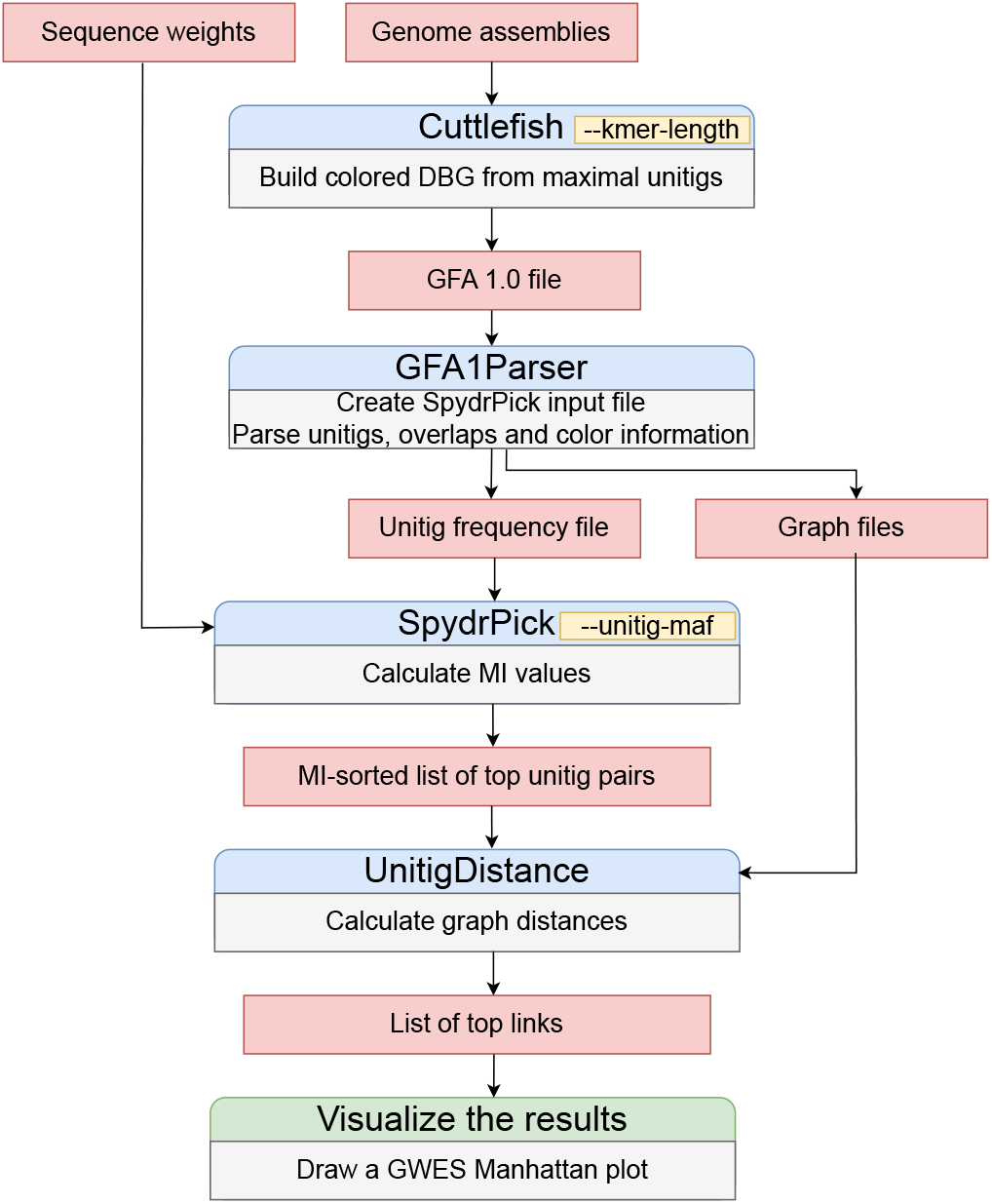
Overview of the PAN-GWES pipeline with the main user-definable parameters highlighted. Assemblies are the only required input, enabling the PAN-GWES to generate phenotype-free biological hypotheses, in a complementary manner to traditional GWAS methods.

The coloured compacted de Bruijn graph is initially built using the computationally efficient Cuttlefish algorithm with a given k-mer length. Linkage disequilibrium between pairs of variants is measured using Mutual Information (MI), with an additional weighting to account for population structure as described previously (3). The MI-based method is specifically designed to detect long-distance pairs that show an elevated association with respect to a background distribution, and thus relies on a distance measure to filter out short-distance pairs.

Without a multiple sequence alignment based on a single reference genome, it is not straight-forward to define a unique distance between variants observed in a population of genomes. To approximate the distance for a pair of unitigs we instead compute the distribution of distances over the de Bruijn graphs containing both of the corresponding unitigs and use the mean of the distribution as an effective measure of their genomic proximity. To avoid calculation of MI for non-informative unitigs, we included options for filtering out unitigs based on their Minor Allele Frequency (MAF) and the inferred genomic distance. The PAN-GWES method leverages the computational efficiencies of the SpydrPick algorithm to rapidly calculate the pairwise MI values of millions of unitigs pairs, allowing it to scale to large and diverse pangenomes such as those of *Escherichia coli*.

A critical component of the PAN-GWES pipeline is the initial choice of k-mer length. Longer k-mers are more specific and can capture fine-scale mutations. Conversely, shorter k-mers allow for greater sequence diversity and are better at capturing differences in gene content (12, 13). Since obtaining a good balance between shorter and longer k-mers depends on the amount of diversity present in the target species, we allow the user to run the PAN-GWES pipeline repeatedly with different values of k to explore and compare the resulting LD patterns.

### Pangenome GWES analysis identifies novel links of co-selection in *Streptococcus pneumoniae*

To verify that our graph-based distance estimates can identify signals of epistasis and co-selection we compared the results from our pangenome approach to the epistatic and co-evolutionary hits identified by running the previously published SpydrPick algorithm on a multiple sequence alignment (MSA) built from a collection of 3042 *Streptococcus pneumoniae* genomes (17). The genomes were sequenced from isolates collected in Maela, a refugee camp on the Thai-Myanmar border between 2007-2010. SpydrPick uses the distance between variants in an MSA, usually built by aligning sequencing reads to a reference genome. This assumes that the distance between sites in the reference is a good proxy for the distance between sites in the full collection of genomes. The Maela collection of *S. pneumoniae* genomes was sequenced in 2010-2011 using short reads 75-bp in length. Consequently, the resulting assemblies are highly fragmented (average contig length of 33,191 bp and average N50 of 65,656 bp) which represents a challenge for our algorithm.

Figure 3, indicates that despite the difficulties caused by fragmentation, our pangenome distance-based approach is able to identify similar signals to the SpyrdPick algorithm albeit with some loss in sensitivity. The strong co-evolutionary signal between the penicillin-binding proteins (PBPs) pbpX and pbp2B was clearly identified using both algorithms (shown in purple). The PBP proteins are targets for betalactam antibiotics and modifications in both of these proteins are the primary resistance mechanism for multiple classes of beta-lactams (18, 19). Similarly, the pangenome-based algorithm identified strong links (yellow) between SPN23F16620 (divIVA) and the gene cluster SPN23F19480-SPN23F19500 which is located directly upstream of the gene encoding the toxin and key virulence factor pneumolysin (ply/SPN23F19470). divIVA encodes a cell morphogenesis factor and these coupling links have been hypothesised to be the consequence of virulence proteins interacting at the cell surface (2, 20).

**Fig. 3.**
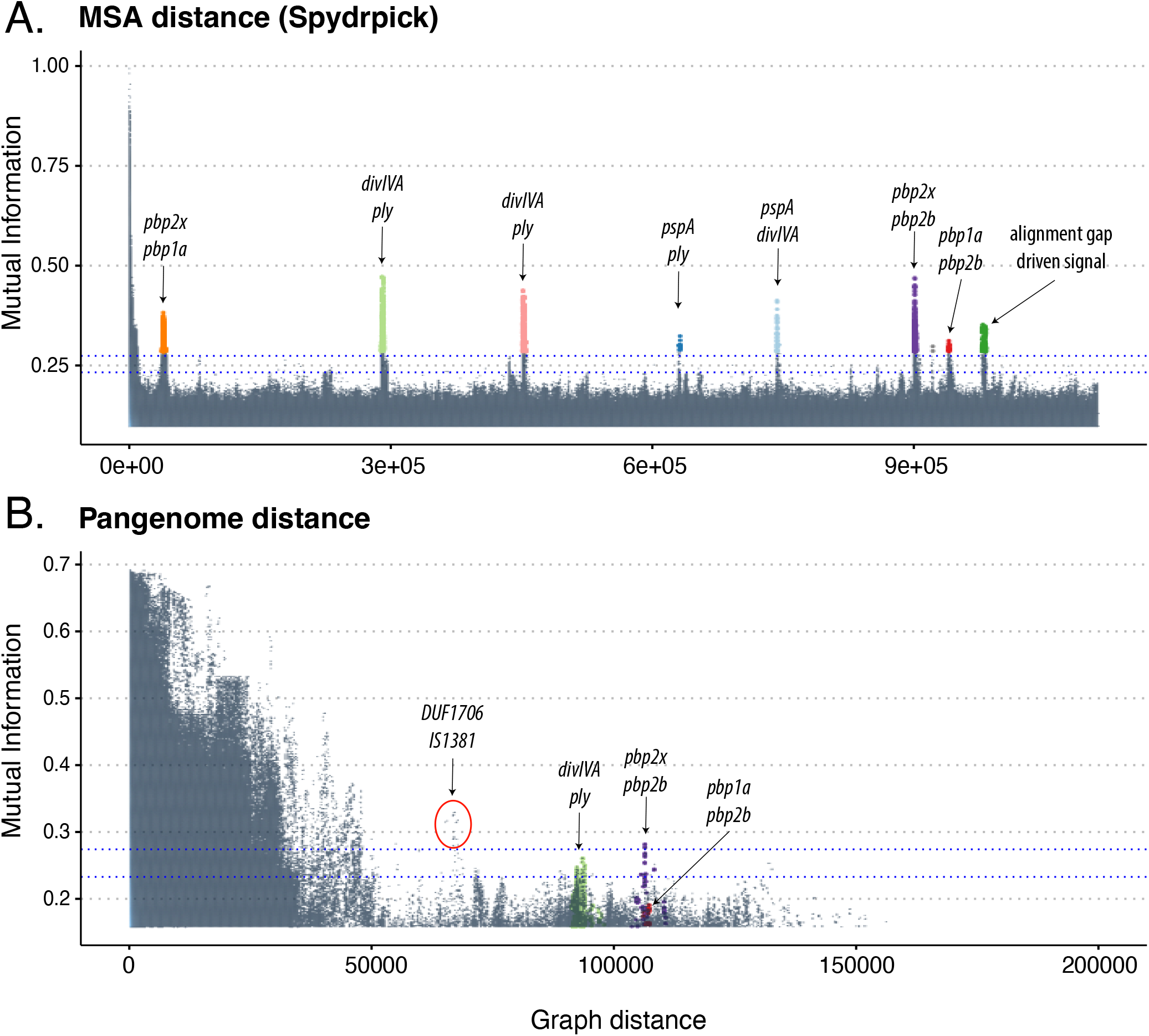
Manhattan plots indicating the strength of linkage disequilibrium, using mutual information, versus the distance separating loci using a single reference and the SpydrPick algorithm (A) and the PAN-GWES algorithm (B). Separate colours are given to each link and are consistent between the plots. Despite the highly fragmented nature of the dataset reducing the sensitivity of the PAN-GWES approach, similar signals of co-selection between the penicillin binding proteins were observed. The strongest link observed in the PAN-GWES approach (red circle) between a DUF1706 domain-containing protein and the putative insertion sequence IS1381 was obscured when relying on a single reference genome as the location of the insertion sequence varies considerably.

In addition to rediscovering known associations, our pangenome graph approach detected a number of novel linked loci that were obscured when relying on only a single reference. This included associations between variants in a ligand binding protein (YpsA) and a Domain of Unknown Function 4231 (DUF4231) containing protein. Expression of DUF4231 containing proteins has been associated with both exposure to cigarette smoke and macrolides (21, 22). A DUF4231 containing protein is also part of the pneumolysis (Ply) operon, encoding a cytolytic pore-forming toxin, and a primary virulence factor for this bacterium (23). Within-host variation in YpsA has been linked with penicillin treatment in the same cohort using a unitig based genome-wide association study (24). This variation was hypothesised to allow pneumococci to reduce their metabolism and cell division, allowing the population to persist in periods of stress when the antibiotic was present. The association between these two loci could be due to their involvement in the same stress response pathways.

The association with the highest MI value for distant unitigs identified using our pangenome approach (red circle) was found between SPN23F08610 (dfsB), a DUF1706 domain-containing protein and the putative insertion sequence IS1381 (SPN23F08630). dfsB is widely distributed across bacterial species and has been associated with inducing cell death in nearby colonies of bacteria (25). The average distance between unitigs from these genes in the de Bruijn graph was 67,000bp. However, the multicopy IS-element is highly mobile and has been observed to insert close (<2000bp) to dfsB in several assemblies including ATCC-700669 (Supplementary Figure 4). The link with an insertion sequence could be associated with phenotypic switching caused by the movement of mobile elements or alternatively it could be an artefact introduced by the highly fragmented assemblies in the Maela collection. To further validate the identified co-evolutionary signal, it is important to apply the PAN-GWES algorithm to enhanced assemblies, potentially using long-read technology.

### Accurate long read assemblies reveal new evidence of co-selection and epistasis in *Enterococcus faecalis*

As the graph-based genomic distance approximation is expected to be most accurate with complete genome assemblies, where longer pangenome distances can be accurately inferred, we tested our method on a large and representatively sampled *E. faecalis* dataset consisting of Illumina and Oxford Nanopore hybrid assemblies (26). In total, we considered 332 complete circular chromosomes, combined with 43 unpublished near-complete chromosomes from the same population, leading to a total of 375 genomes. Figure 4A and Supplementary Figure 5 illustrate the LD-decay pattern based on specific choices of k-mer length and filtering criteria. Longer k-mers are more specific, resulting in larger distances within the pangenome graph. Shorter k-mers enable the identification of associations between more diverse features including gene gain and loss. The stability of the LD signals as a function of approximated genomic distance demonstrates the attractive features of long-read based assemblies as an input to a pangenomic GWES analysis. Moreover, a comparison of chromosome distances calculated using a single genome and the PAN-GWES method were highly correlated, indicating the algorithm provides an informative measure of distance (Figure 4B).

**Fig. 4.**
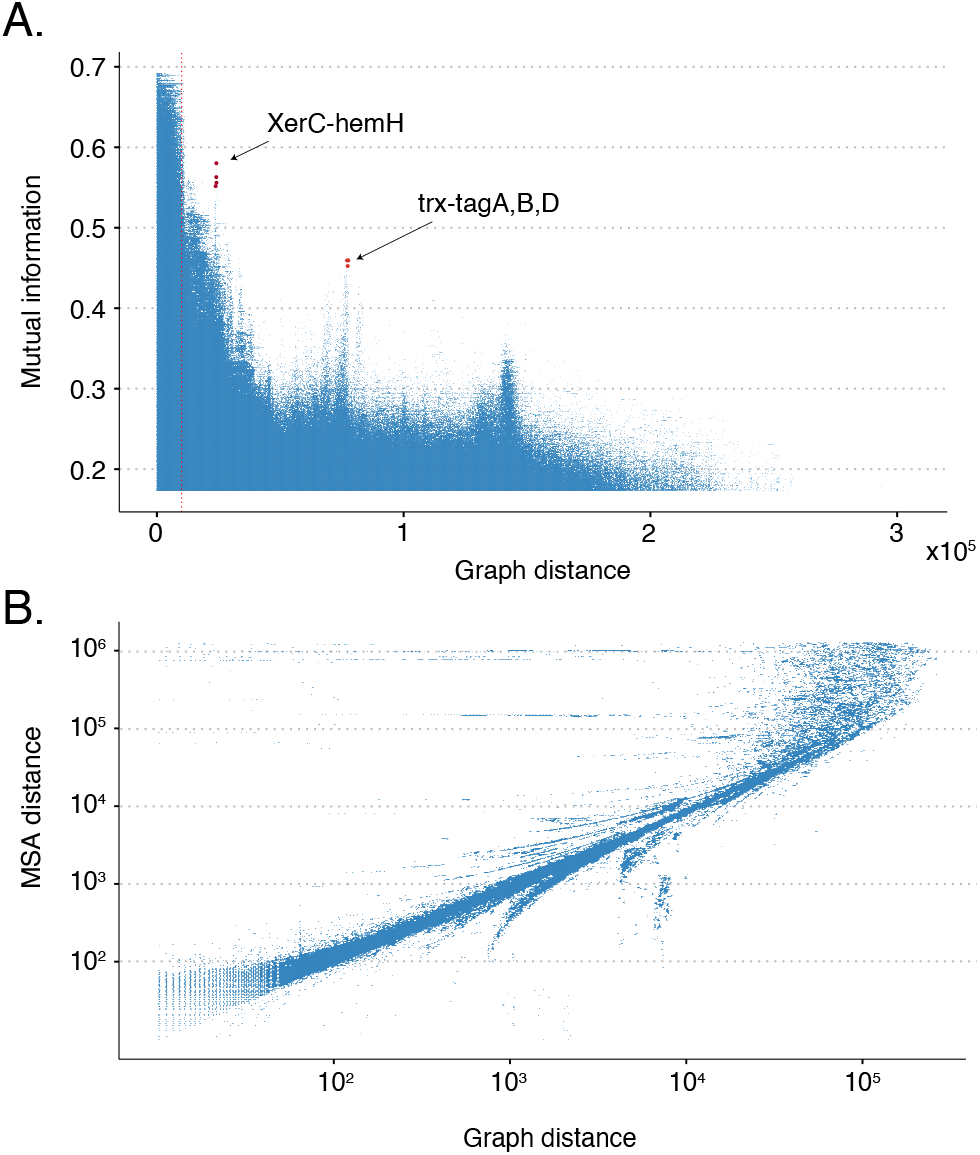
A) A Manhattan plot indicating the strength of linkage disequilibrium measured using mutual information versus the genomic distance in the graph. Strong links of co-selection were identified between an intergenic region adjacent to the trxB gene and the three genes tagA, B and D in addition to a link between a site-specific XerC family tyrosine recombinase and an intergenic region between a tRNAs gene and a ferrochelatase-encoding hemH. B) Pairwise distances between loci within a single genome and those found using the PAN-GWES graph based approach. The correlation, particularly over shorter distances, indicates that the PAN-GWES algorithm can accurately distinguish LD driven by proximity within the genome from LD indicative of co-selection or epistasis. The horizontal lines are caused by k-mers associated with mobile elements that appear in different locations within each genome but are fixed within the single reference.

Screening of the top hits within the *E. faecalis* collection resulted in 66 unitig pairs, further filtered to 29 unique pairs when considering individual genes and their intergenic regions (Supplementary Table 1). Of these, intergenic regions were overrepresented, as 65.5% (19/29) of hit pairs exhibited an intergenic region in at least one of the pairs of unitigs. To assess whether any signals were associated with the clinical setting, the presence of top hits was compared between isolates from hospital and non-hospital sources, with the over-representation of intergenic hits also reflected in the hospital-associated isolates. Five unique hits were more prevalent in *E. faecalis* isolates from hospitalised patients compared to other sources (Supplementary Table 1), four of which included the intergenic region between a hypothetical protein and a thioredoxin reductase-encoding trxB (Chi-Square test, p < 0.05). TrxB is a conserved detoxifying enzyme and part of the thioredoxin complex, globally involved in the oxidative stress response (27, 28). In addition to another hypothetical protein, the trxB intergenic hit was separately linked to three genes (tagA, tagB and tagD) from the cell wall teichoic acid synthesis pathway, of which intact tagB has been indicated a role in evading the complement system activation in a host by modification of peptidoglycan (29). Oxidative stress, both in the environment and during intracellular host infection, as well as the hurdles of the host immune system are both conditions that *E. faecalis* would frequently face in the hospital setting, potentially explaining the multiple links between the two functions. Another hit enriched in hospital isolates, was identified between a site-specific XerC family tyrosine recombinase and an intergenic region between a tRNA gene and a ferrochelatase-encoding hemH which adds iron to heme. This link could be explained by the connection of the integrase gene of a phage to its integration site, perhaps more so than directly to the ferrochelatase. As intergenic regions often harbour regulatory machinery of the nearby genes, their overrepresentation within the top hits demonstrate the potential of our phenotype-free approach to uncover regulatory signals that can be important in pathogenesis and which usually would be detectable from population transcriptomic data only (30).

## Discussion

The decreasing cost and improved scalability of both short- and long-read sequencing are continuing to rapidly increase the availability of high-quality population genomics data for many bacterial species, in particular for those with relevance to public health. Currently there is untapped potential in using these data to study bacterial evolution and adaptation. Genome-wide epistasis and co-selection study (GWES) is a recently emerged tool for uncovering drivers of change in genomes through LD pattern analysis without assuming availability of phenotypic data from population-wide screening. However, currently these approaches are limited to consider only the core genome. To fill this gap, we introduced a PAN-GWES approach that allows for a more holistic view over the population patterns of genomic co-variation.

We identified strong associations between known antibiotic resistance genes in *S. pneumoniae* in addition to links enriched in hospital derived *E. faecalis* isolates, which demonstrate the potential of our approach to identify genes and regions within the genome that could be the target of future studies or interventions aimed at reducing the burden of infectious disease. A key innovation in our method is the use of a coloured and compacted de Bruijn graphs to obtain a computationally scalable measure of the genomic distance between loci within a species pangenome.

While highly fragmented assemblies represent a challenge for our algorithm, we demonstrate that although it can lead to a reduction in sensitivity over reference based approaches, we are nevertheless able to identify signals of epistasis and co-selection that would be missed by previous approaches. Notably, the ‘missing’ signal identified in our analysis of the pneumococcal pangenome included genes previously associated with antimicrobial treatment. This further demonstrates the ability of our method to detect clinically important associations without requiring phenotypic meta-data. When dealing with highly fragmented data sets we recommend the use of both reference based methods, such as SpydrPick, and PAN-GWES to take advantage of the strengths of both tools. As advancements in long-read technologies continue to deliver increased precision and affordability, we anticipate that genome assemblies will shift towards completeness or near-completeness as the standard. Consequently, the efficacy of our algorithm is likely to improve, providing more accurate inferences on contemporary genome datasets, as exemplified in our *E. faecalis* analysis.

## Methods

### de Bruijn graph construction

Let *S* = (*S*_1_, …, *S*_*N*_) be a set of N assembled DNA sequence strings over the DNA alphabet Σ = {*A, C, G, T*}. A substring of length *k* that is contained within a sequence in *S* is called a k-mer. Given a k-mer *s*, let 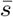 denote the reverse complement of *s*, which is formed by reversing the string and interchanging A and T and interchanging C and G. We denote by *ŝ* the lexicographically smaller string between *s* and 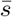, which is called the canonical k-mer. Using canonical k-mers allows us to account for each location in a genome once whilst accounting for both reverse complementary sequences. For a k-mer *s*, pre(*s, l*) and suf(*s, l*) denote the prefix and the suffix of length *l* of *s*, respectively.

Given a set of strings *S* and an odd integer *k >* 0, we define the de Bruijn graph as a bidirected multigraph *G*_*S,k*_ = (*V, E*) where the set of vertices *V* is exactly the set of canonical k-mers contained in the sequences in *S*. Two vertices *v* and *w* in *V* are connected by an edge *e* if and only if there exists a (*k* + 1)-mer *z* in *S* such that pre 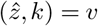 and suf 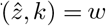. The choice of an odd k-mer length ensures that a k-mer can never be its own reverse complement, which would cause ambiguity in the graph.

A sequence of distinct vertices is called a path if every two adjacent vertices in the path are connected by an edge in G. The path *p* = (*v*_1_, …, *v*_*m*_) is called non-branching if (1) each internal vertex *v*_*i*_, *i* = 2, …, *m*− 1, has exactly two edges that connect to different sides of *v*_*i*_ and (2) the first and last vertices *v*_1_ and *v*_*m*_ have exactly one edge on the side which connects to *v*_2_ and *v*_*m*−1_ respectively. A non-branching path is said to be maximal if the path cannot be extended by a vertex on either side without branching. For convenience, maximal non-branching paths are reduced to a single vertex, referred to as a ‘unitig’, leading to a compacted de Bruijn graph (cdBG).

### Coloured de Bruijn graph

We can extend the cdBG to include colours representing which k-mer is present in which genome. A coloured de Bruijn graph is a graph G=(V,E,C) in which (V,E) is a dBG and C is a set of colours such that each vertex *v* ∈ *V* maps to a subset of *C*. A path through this graph is called colour-coherent if all its vertices share the same colour set. We define the coloured compacted de Bruijn graph as a bidirected multigraph *G*_*S,k*_ = (*V, E, C*), which is constructed by collapsing all of its maximal colour-coherent non-branching paths into single vertices, which are called unitigs.

### Genomic distance calculation

In *G*_*S,k*_, the length of a path is simply the number of edges in the path. However, a path *p* = (*v*_1_, …, *v*_*m*_) in the compacted de Bruijn graph obtained from *G*_*S,k*_ consists of unitigs which in itself are paths in *G*_*S,k*_. Therefore, to measure the length of *p*, special attention must be given to the actual number of edges traversed with respect to the uncompacted graph. We define the shortest distance between two unitigs *p* and *q* in *V* to be the length of the path starting from *p* and ending at *q*, traversing via edges in the uncompacted graph, with minimal length. The path is calculated using a parallel version of Dijkstra’s algorithm (31). To define the distance between unitigs in a compact coloured de Bruijn graph, we calculate the distance between *p* and *q* for each subgraph *v* ∈ *V* where *v* maps to a single colour in *C*. The average distance over all colours is then taken.

### Implementation

The PAN-GWES program is an open-source program implemented in C++ that handles the efficient construction of the colour-induced de Bruijn subgraphs and the following shortest path distance calculations. It supports parallel execution and further improves run time by smart arrangement of the graph search jobs. The program also incorporates a convenient parser for Graphical Fragment Assembly 1.0 (GFA1) formatted files in order to construct all the necessary files used in the full pipeline.

The full PAN-GWES pipeline consists of four stages. First, Cuttlefish is utilised to construct the coloured compacted de Bruijn graph in GFA1 Format which is then parsed by PAN-GWES to prepare various graph data files with colour information and a unitig frequency file to be used as an input file for SpydrPick (3, 15). Next, the list of top candidate unitig pairs is calculated with the SpydrPick algorithm. Finally, the PAN-GWES program is used to calculate the distances between unitigs in the subgraph containing the nodes present in a single genome within the full coloured de Bruijn.

The PAN-GWES program is available under the MIT License from GitHub: https://github.com/jurikuronen/PANGWES.

### Sequencing, assembly and variant calling

All the circular chromosome sequences of the fully contiguous hybrid assemblies (n=332) were included from the previously collated *E. faecalis* dataset (26). The collection was supplemented with 43 new, partially contiguous *E. faecalis* hybrid assemblies, constructed from Illumina short-read and Oxford Nanopore Technologies (ONT) long-read sequences using a hybrid assembly pipeline (https://github.com/arredondo23/hybrid_assembly_slurm) with Unicycler v.0.4.7 (32). The newly introduced isolates were delineated into clusters using PopPUNK (12) v.1.1.5 with the – assign-query mode against the previously-curated *E. faecalis* database and clustering scheme (26). The unique pan-GWES top hits were annotated using annotate_hits_pyseer of the pyseer tool v.1.3.9 (6) against selected hybrid assemblies from the collection serving as references to cover all of the top hits. Unitig-caller v.1.3.0 (7, 9) with –simple mode was used for creating presence/absence matrix of the top hit pairs across the *E. faecalis* collection. Annotated hits were manually curated and hit pairs allocated with the same annotations were merged. To test the significance of differences in proportions of each PAN-GWES top hit pair between hospital- and non-hospital-associated *E. faecalis* isolates, two-sided Fisher’s exact test was used when there were any counts less than five and Pearson’s Chi-squared test with higher counts than five. Both tests were performed with a significance threshold of p < 0.05.

## Data and materials availability

The Nanopore sequencing data for the 43 *E. faecalis* isolates are available in the European Nucleotide Archive (ENA) under the accession number PRJEB40976. The sequencing data for the remaining *E. faecalis* isolates is available under the original study accession number PRJEB28327. The raw pneumococcal sequencing reads are available from the NCBI Sequencing Read Archive (SRA) under the original study accession numbers ERP000435, ERP000483, ERP000485, ERP000487, ERP000598, ERP000599.

## Supplementary Figures

**Supplementary Figure 1.**
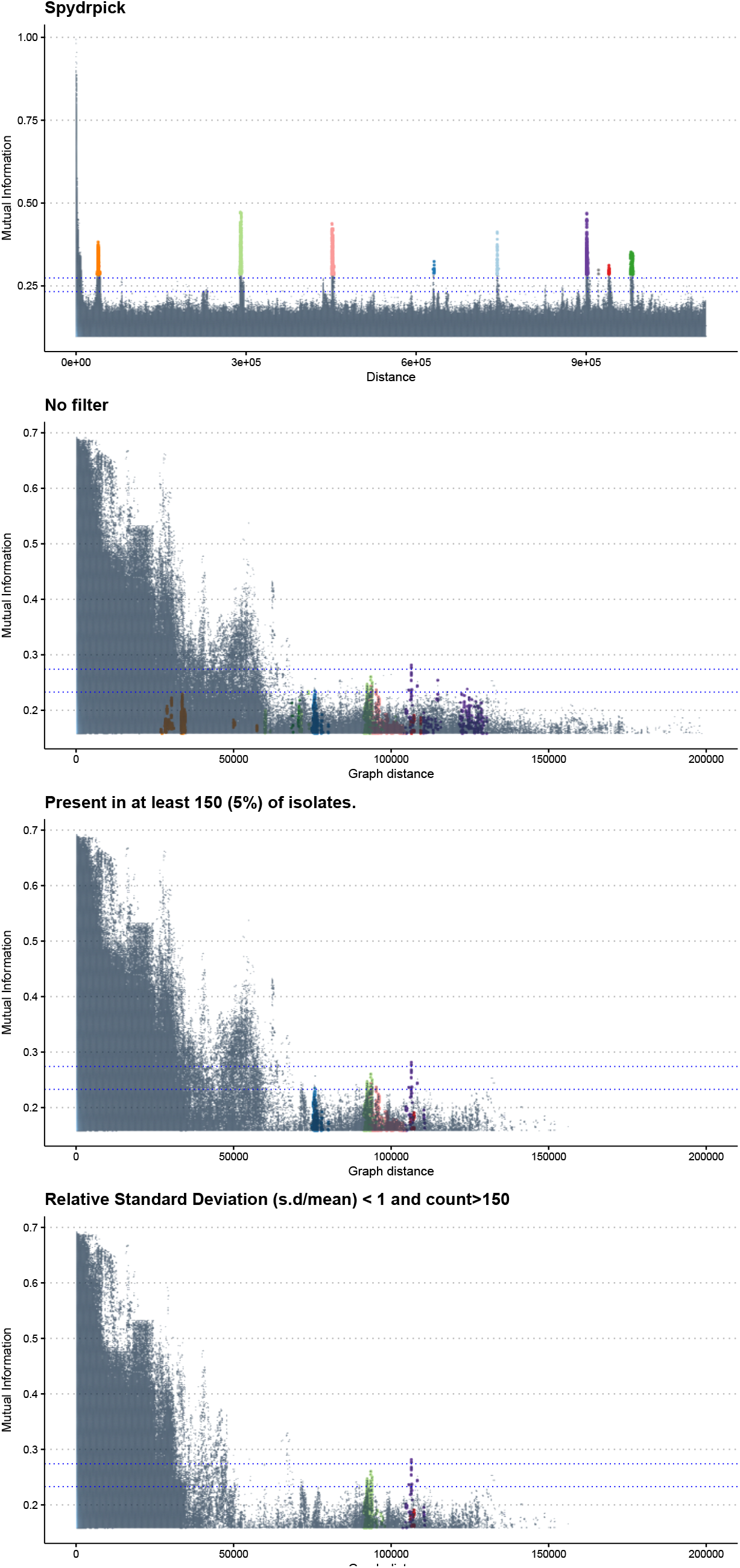
Manhattan plots indicating the distribution of MI values by distance with different filters in the PAN-GWES pipeline applied to the *S. pneumoniae* dataset. The top facet indicates the results of the Spydrpick algorithm that relies on a reference genome alignment.

**Supplementary Figure 2.**
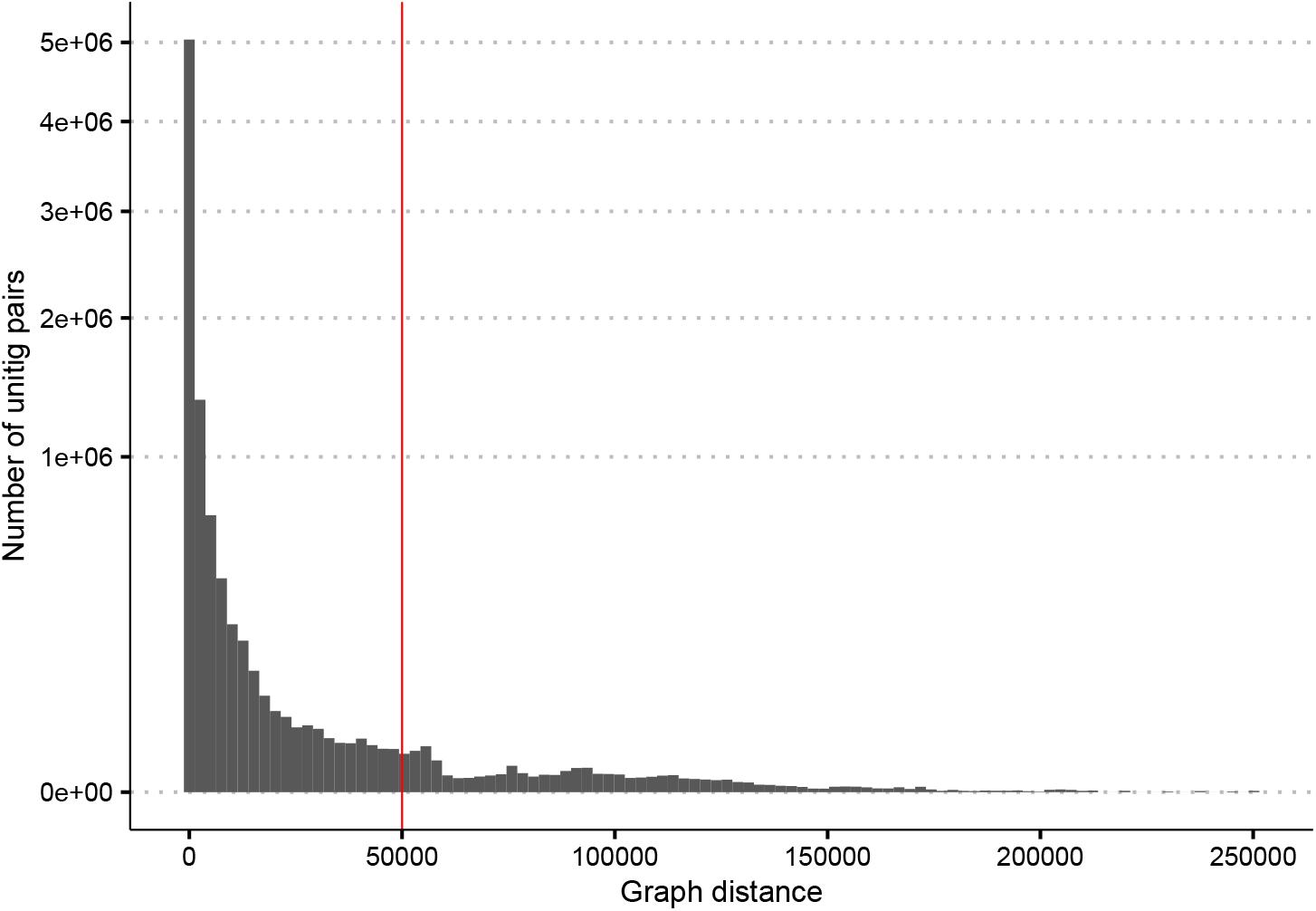
A histogram indicating the number of unitig pairs versus the average distance across each colour in the coloured de-Bruijn graph for the pneumococcal dataset. Only pairs above 50,000bp were considered in the final analysis.

**Supplementary Figure 3.**
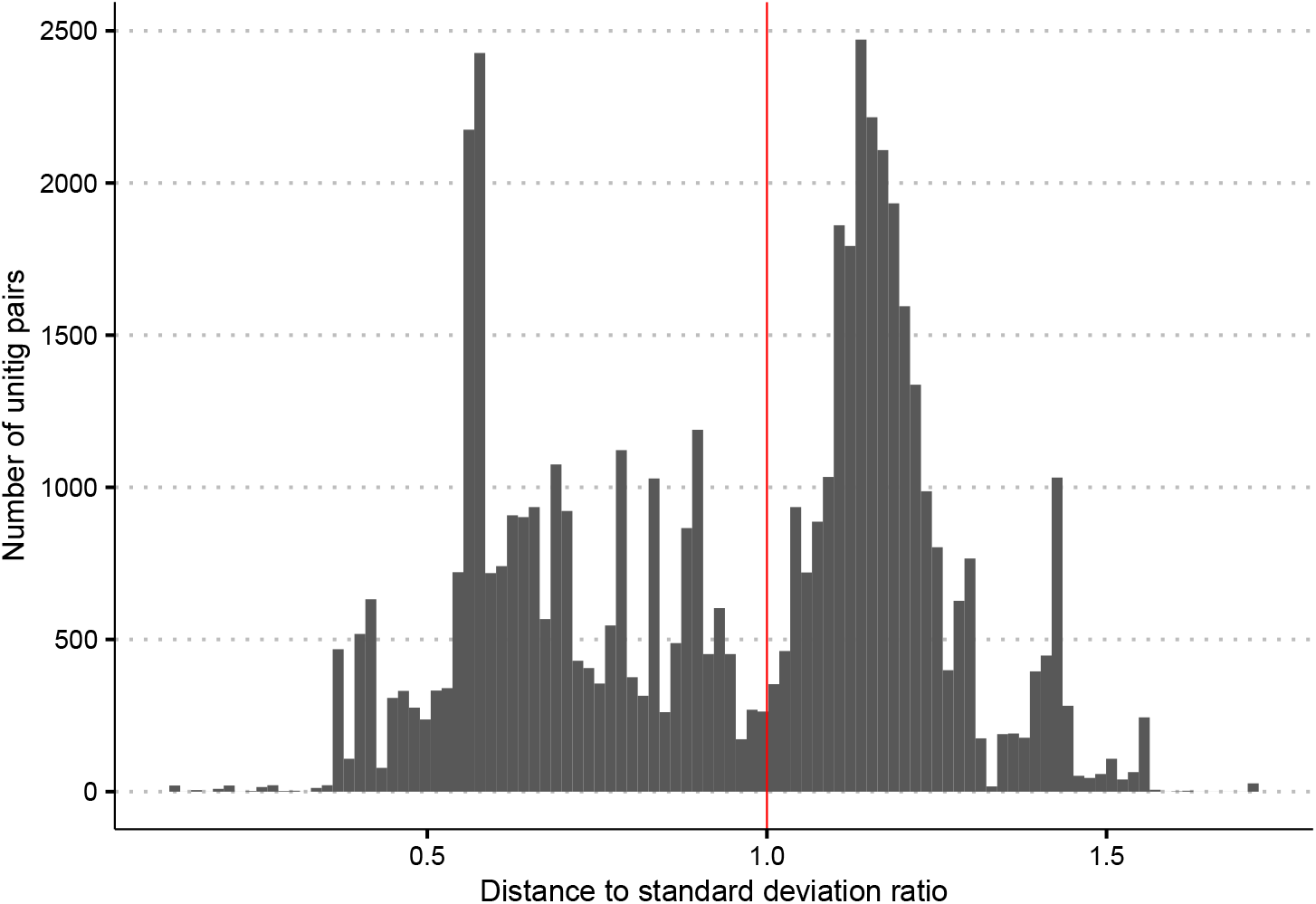
The number of unitig pairs versus the standard deviation of their distance in each subgraph represented by an individual colour in the full coloured de Bruijn graph.

**Supplementary Figure 4.**
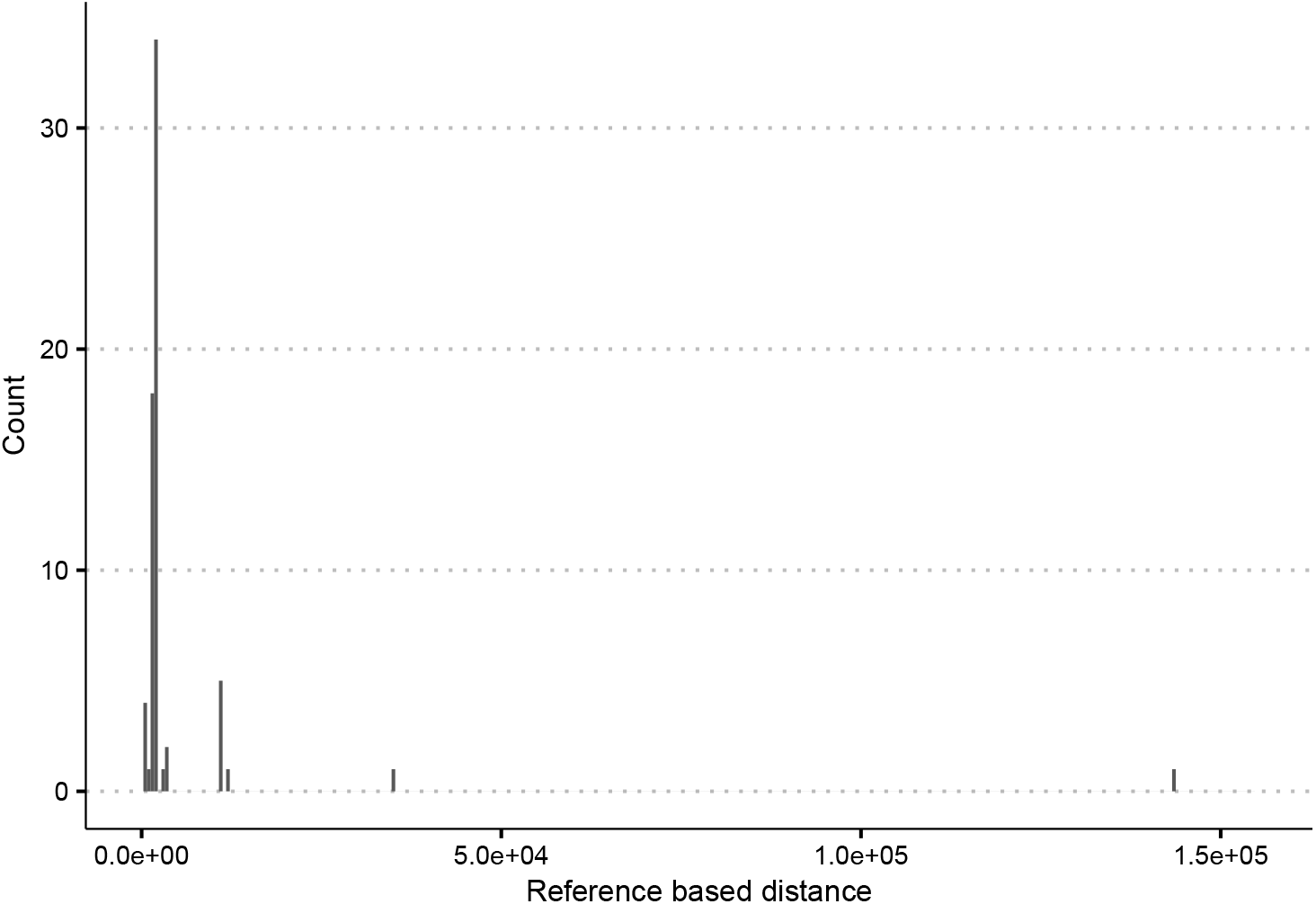
A histogram of the distances between two unitigs linked to SPN23F08610 (dfsB), a DUF1706 domain-containing protein and the putative insertion sequence IS1381 (SPN23F08630) when they appeared on the same contig of an assembly. The cluster of short distances indicates that this link may be impacted by the highly fragmented assemblies in this dataset and highlights the importance of subsequent laboratory experiments to confirm identified associations.

**Supplementary Figure 5.**
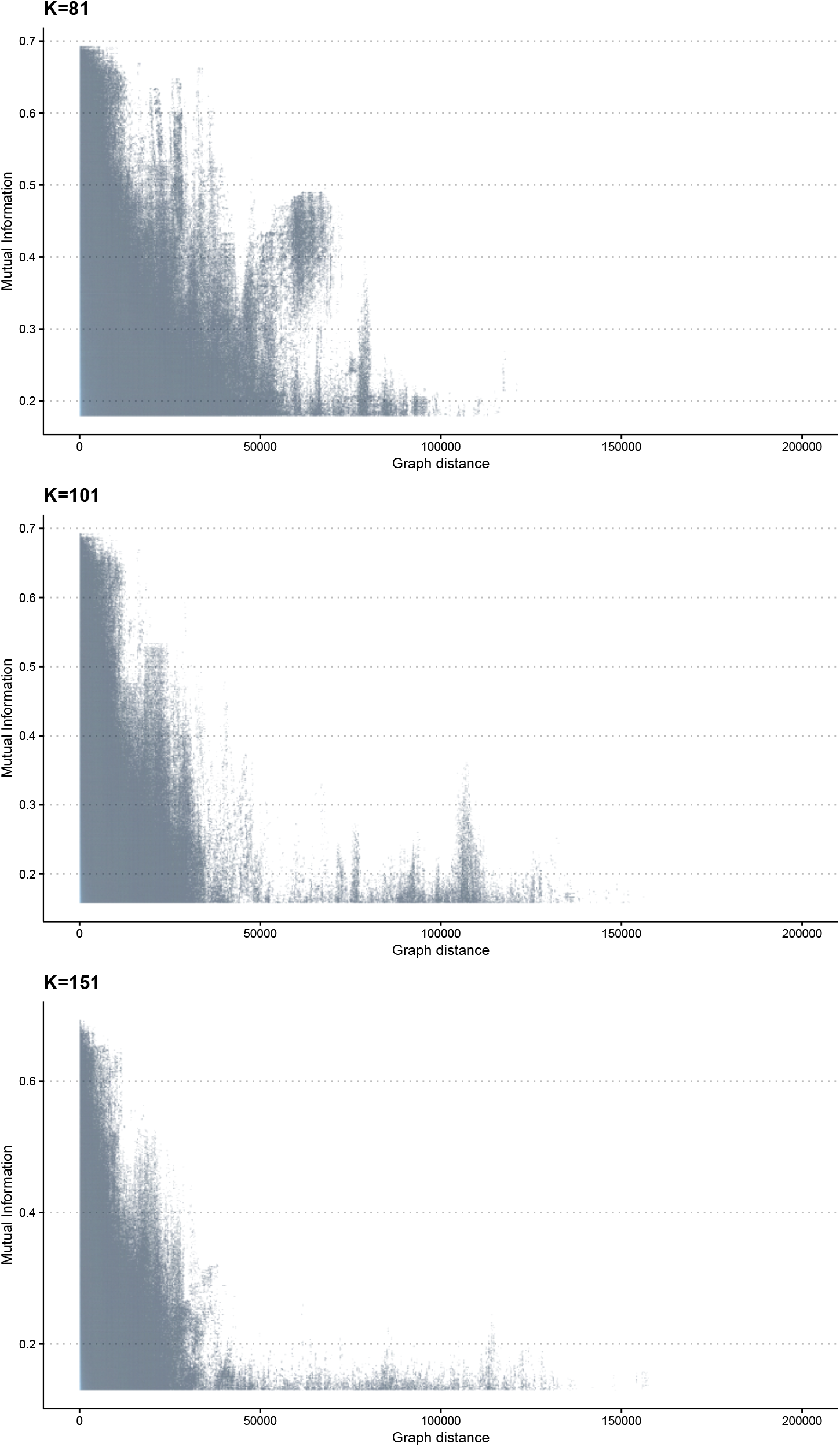
Manhattan plots indicating the distribution of MI values by distance with different k-mer lengths in the PAN-GWES pipeline applied to the *E. faecalis* dataset.

## References

1. Brian J Arnold, Michael U Gutmann, Yonatan H Grad, Samuel K Sheppard, Jukka Corander, Marc Lipsitch, and William P Hanage. Weak epistasis may drive adaptation in recombining bacteria. Genetics, 208(3):1247–1260, March 2018. ISSN 0016-6731. doi: 10.1534/genetics.117.300662.

2. Marcin J Skwark, Nicholas J Croucher, Santeri Puranen, Claire Chewapreecha, Maiju Pesonen, Ying Ying Xu, Paul Turner, Simon R Harris, Stephen B Beres, James M Musser, Julian Parkhill, Stephen D Bentley, Erik Aurell, and Jukka Corander. Interacting networks of resistance, virulence and core machinery genes identified by genome-wide epistasis analysis. PLoS Genet., 13(2):e1006508, February 2017. ISSN 1553-7390, 1553-7404. doi: 10.1371/journal.pgen.1006508.

3. Johan Pensar, Santeri Puranen, Brian Arnold, Neil MacAlasdair, Juri Kuronen, Gerry Tonkin-Hill, Maiju Pesonen, Yingying Xu, Aleksi Sipola, Leonor Sánchez-Busó, John A Lees, Claire Chewapreecha, Stephen D Bentley, Simon R Harris, Julian Parkhill, Nicholas J Croucher, and Jukka Corander. Genome-wide epistasis and co-selection study using mutual information. Nucleic Acids Res., 47(18):e112, October 2019. ISSN 0305-1048, 1362-4962. doi: 10.1093/nar/gkz656

4. Janetta Top, Sergio Arredondo-Alonso, Anita C Schürch, Santeri Puranen, Maiju Pesonen, Johan Pensar, Rob J L Willems, and Jukka Corander. Genomic rearrangements uncovered by genome-wide co-evolution analysis of a major nosocomial pathogen, enterococcus faecium. Microb Genom, 6(12), December 2020. ISSN 2057-5858. doi: 10.1099/mgen.0.000488.

5. Claire Chewapreecha, Johan Pensar, Supaksorn Chattagul, Maiju Pesonen, Apiwat Sangphukieo, Phumrapee Boonklang, Chotima Potisap, Sirikamon Koosakulnirand, Edward J Feil, Susanna Dunachie, Narisara Chantratita, Direk Limmathurotsakul, Sharon J Peacock, Nick P J Day, Julian Parkhill, Nicholas R Thomson, Rasana W Sermswan, and Jukka Corander. Co-evolutionary signals identify burkholderia pseudomallei survival strategies in a hostile environment. Mol. Biol. Evol., 39(1):msab306, October 2021. ISSN 0737-4038. doi: 10.1093/molbev/msab306.

6. John A Lees, Marco Galardini, Stephen D Bentley, Jeffrey N Weiser, and Jukka Corander. pyseer: a comprehensive tool for microbial pangenome-wide association studies. Bioinformatics, 34(24):4310–4312, December 2018. ISSN 1367-4803, 1367-4811. doi: 10.1093/bioinformatics/bty539.

7. John A Lees, T Tien Mai, Marco Galardini, Nicole E Wheeler, Samuel T Horsfield, Julian Parkhill, and Jukka Corander. Improved prediction of bacterial Genotype-Phenotype associations using interpretable Pangenome-Spanning regressions. MBio, 11(4), July 2020. ISSN 2150-7511. doi: 10.1128/mBio.01344-20.

8. Sarah G Earle, Chieh-Hsi Wu, Jane Charlesworth, Nicole Stoesser, N Claire Gordon, Timothy M Walker, Chris C A Spencer, Zamin Iqbal, David A Clifton, Katie L Hopkins, Neil Woodford, E Grace Smith, Nazir Ismail, Martin J Llewelyn, Tim E Peto, Derrick W Crook, Gil McVean, A Sarah Walker, and Daniel J Wilson. Identifying lineage effects when controlling for population structure improves power in bacterial association studies. Nat Microbiol, 1: 16041, April 2016. ISSN 2058-5276. doi: 10.1038/nmicrobiol.2016.41.

9. Guillaume Holley and Páll Melsted. Bifrost: highly parallel construction and indexing of colored and compacted de bruijn graphs. Genome Biol., 21(1):249, September 2020. ISSN 1465-6906. doi: 10.1186/s13059-020-02135-8.

10. Anton Bankevich, Sergey Nurk, Dmitry Antipov, Alexey A Gurevich, Mikhail Dvorkin, Alexander S Kulikov, Valery M Lesin, Sergey I Nikolenko, Son Pham, Andrey D Prjibelski, Alexey V Pyshkin, Alexander V Sirotkin, Nikolay Vyahhi, Glenn Tesler, Max A Alekseyev, and Pavel A Pevzner. SPAdes: a new genome assembly algorithm and its applications to single-cell sequencing. J. Comput. Biol., 19(5):455–477, May 2012. ISSN 1066-5277, 1557-8666. doi: 10.1089/cmb.2012.0021.

11. Rachel M Colquhoun, Michael B Hall, Leandro Lima, Leah W Roberts, Kerri M Malone, Martin Hunt, Brice Letcher, Jane Hawkey, Sophie George, Louise Pankhurst, and Zamin Iqbal. Pandora: nucleotide-resolution bacterial pan-genomics with reference graphs. Genome Biol., 22(1):267, September 2021. ISSN 1465-6906. doi: 10.1186/s13059-021-02473-1.

12. John A Lees, Simon R Harris, Gerry Tonkin-Hill, Rebecca A Gladstone, Stephanie W Lo, Jeffrey N Weiser, Jukka Corander, Stephen D Bentley, and Nicholas J Croucher. Fast and flexible bacterial genomic epidemiology with PopPUNK. Genome Res., 29(2):304–316, February 2019. ISSN 1088-9051, 1549-5469. doi: 10.1101/gr.241455.118.

13. Magali Jaillard, Leandro Lima, Maud Tournoud, Pierre Mahé, Alex van Belkum, Vincent Lacroix, and Laurent Jacob. A fast and agnostic method for bacterial genome-wide association studies: Bridging the gap between k-mers and genetic events. PLoS Genet., 14 (11):e1007758, November 2018. ISSN 1553-7390, 1553-7404. doi: 10.1371/journal.pgen.1007758.

14. Zamin Iqbal, Mario Caccamo, Isaac Turner, Paul Flicek, and Gil McVean. De novo assembly and genotyping of variants using colored de bruijn graphs. Nat. Genet., 44(2):226–232, January 2012. ISSN 1061-4036, 1546-1718. doi: 10.1038/ng.1028.

15. Jamshed Khan and Rob Patro. Cuttlefish: fast, parallel and low-memory compaction of de bruijn graphs from large-scale genome collections. Bioinformatics, 37(Suppl_1):i177–i186, July 2021. ISSN 1367-4803, 1367-4811. doi: 10.1093/bioinformatics/btab309.

16. Andrea Cracco and Alexandru I Tomescu. Extremely fast construction and querying of compacted and colored de bruijn graphs with GGCAT. Genome Res., May 2023. ISSN 1088-9051, 1549-5469. doi: 10.1101/gr.277615.122.

17. Claire Chewapreecha, Simon R Harris, Nicholas J Croucher, Claudia Turner, Pekka Marttinen, Lu Cheng, Alberto Pessia, David M Aanensen, Alison E Mather, Andrew J Page, Susannah J Salter, David Harris, Francois Nosten, David Goldblatt, Jukka Corander, Julian Parkhill, Paul Turner, and Stephen D Bentley. Dense genomic sampling identifies highways of pneumococcal recombination. Nat. Genet., 46(3):305–309, March 2014. ISSN 1061-4036, 1546-1718. doi: 10.1038/ng.2895.

18. B G Spratt. Resistance to antibiotics mediated by target alterations. Science, 264(5157): 388–393, April 1994. ISSN 0036-8075. doi: 10.1126/science.8153626.

19. T Grebe and R Hakenbeck. Penicillin-binding proteins 2b and 2x of streptococcus pneumoniae are primary resistance determinants for different classes of beta-lactam antibiotics. Antimicrob. Agents Chemother., 40(4):829–834, April 1996. ISSN 0066-4804. doi: 10.1128/AAC.40.4.829.

20. Aurore Fleurie, Sylvie Manuse, Chao Zhao, Nathalie Campo, Caroline Cluzel, Jean-Pierre Lavergne, Céline Freton, Christophe Combet, Sébastien Guiral, Boumediene Soufi, Boris Macek, Erkin Kuru, Michael S VanNieuwenhze, Yves V Brun, Anne-Marie Di Guilmi, Jean-Pierre Claverys, Anne Galinier, and Christophe Grangeasse. Interplay of the serine/threonine-kinase StkP and the paralogs DivIVA and GpsB in pneumococcal cell elongation and division. PLoS Genet., 10(4):e1004275, April 2014. ISSN 1553-7390, 1553-7404. doi: 10.1371/journal.pgen.1004275.

21. Sam Manna, Alicia Waring, Angelica Papanicolaou, Nathan E Hall, Steven Bozinovski, Eileen M Dunne, and Catherine Satzke. The transcriptomic response of streptococcus pneumoniae following exposure to cigarette smoke extract. Sci. Rep., 8(1):15716, October 2018. ISSN 2045-2322. doi: 10.1038/s41598-018-34103-5.

22. Blue Goad and Laura K Harris. Identification and prioritization of macrolideresistance genes with hypothetical annotation instreptococcus pneumoniae. Bioinformation, 14(9):488–498, December 2018. ISSN 0973-2063. doi: 10.6026/97320630014488.

23. Emily J Stevens, Daniel J Morse, Dora Bonini, Seána Duggan, Tarcisio Brignoli, Mario Recker, John A Lees, Nicholas J Croucher, Stephen Bentley, Daniel J Wilson, Sarah G Earle, Robert Dixon, Angela Nobbs, Howard Jenkinson, Tim van Opijnen, Derek Thibault, Oliver J Wilkinson, Mark S Dillingham, Simon Carlile, Rachel M McLoughlin, and Ruth C Massey. Targeted control of pneumolysin production by a mobile genetic element in streptococcus pneumoniae. MicrobGenom, 8(4), April 2022. ISSN 2057-5858. doi: 10.1099/mgen.0.000784.

24. Gerry Tonkin-Hill, Clare Ling, Chrispin Chaguza, Susannah J Salter, Pattaraporn Hinfonthong, Elissavet Nikolaou, Natalie Tate, Andrzej Pastusiak, Claudia Turner, Claire Chewapreecha, Simon D W Frost, Jukka Corander, Nicholas J Croucher, Paul Turner, and Stephen D Bentley. Pneumococcal within-host diversity during colonization, transmission and treatment. Nat Microbiol, 7(11):1791–1804, November 2022. ISSN 2058-5276. doi: 10.1038/s41564-022-01238-1.

25. Jonathan D Taylor, Gabrielle Taylor, Stephen A Hare, and Steve J Matthews. Structures of the DfsB protein family suggest a cationic, helical sibling lethal factor peptide. J. Mol. Biol., 428(3):554–560, February 2016. ISSN 0022-2836, 1089-8638. doi: 10.1016/j.jmb.2016.01.013.

26. Anna K Pöntinen, Janetta Top, Sergio Arredondo-Alonso, Gerry Tonkin-Hill, Ana R Freitas, Carla Novais, Rebecca A Gladstone, Maiju Pesonen, Rodrigo Meneses, Henri Pesonen, John A Lees, Dorota Jamrozy, Stephen D Bentley, Val F Lanza, Carmen Torres, Luisa Peixe, Teresa M Coque, Julian Parkhill, Anita C Schürch, Rob J L Willems, and Jukka Corander. Apparent nosocomial adaptation of enterococcus faecalis predates the modern hospital era. Nat. Commun., 12(1):1523, March 2021. ISSN 2041-1723. doi: 10.1038/s41467-021-21749-5.

27. Jessica K Kajfasz, Jorge E Mendoza, Anthony O Gaca, James H Miller, Kristy A Koselny, Marcia Giambiagi-Demarval, Melanie Wellington, Jacqueline Abranches, and José A Lemos. The spx regulator modulates stress responses and virulence in enterococcus faecalis. Infect. Immun., 80(7):2265–2275, July 2012. ISSN 0019-9567, 1098-5522. doi: 10.1128/IAI.00026-12.

28. Tanja Zeller and Gabriele Klug. Thioredoxins in bacteria: functions in oxidative stress response and regulation of thioredoxin genes. Naturwissenschaften, 93(6):259–266, June 2006. ISSN 0028-1042. doi: 10.1007/s00114-006-0106-1.

29. Stefan Geiss-Liebisch, Suzan H M Rooijakkers, Agnieszka Beczala, Patricia Sanchez-Carballo, Karolina Kruszynska, Christian Repp, Tuerkan Sakinc, Evgeny Vinogradov, Otto Holst, Johannes Huebner, and Christian Theilacker. Secondary cell wall polymers of enterococcus faecalis are critical for resistance to complement activation via mannose-binding lectin. J. Biol. Chem., 287(45):37769–37777, November 2012. ISSN 0021-9258, 1083-351X1. doi: 10.1074/jbc.M112.358283.

30. Priyanka Kachroo, Jesus M Eraso, Stephen B Beres, Randall J Olsen, Luchang Zhu, Waleed Nasser, Paul E Bernard, Concepcion C Cantu, Matthew Ojeda Saavedra, María José Arredondo, Benjamin Strope, Hackwon Do, Muthiah Kumaraswami, Jaana Vuopio, Kirsi Gröndahl-Yli-Hannuksela, Karl G Kristinsson, Magnus Gottfredsson, Maiju Pesonen, Johan Pensar, Emily R Davenport, Andrew G Clark, Jukka Corander, Dominique A Caugant, Shahin Gaini, Marita Debess Magnussen, Samantha L Kubiak, Hoang A T Nguyen, S Wesley Long, Adeline R Porter, Frank R DeLeo, and James M Musser. Integrated analysis of population genomics, transcriptomics and virulence provides novel insights into streptococcus pyogenes pathogenesis. Nat. Genet., 51(3):548–559, March 2019. ISSN 1061-4036, 1546-1718. doi: 10.1038/s41588-018-0343-1.

31. E W Dijkstra. A note on two problems in connexion with graphs. Numer. Math., 1(1):269–271, December 1959. ISSN 0029-599X, 0945-3245. doi: 10.1007/BF01386390.

32. Ryan R Wick, Louise M Judd, Claire L Gorrie, and Kathryn E Holt. Unicycler: Resolving bacterial genome assemblies from short and long sequencing reads. PLoS Comput. Biol., 13 (6):e1005595, June 2017. ISSN 1553-734X, 1553-7358. doi: 10.1371/journal.pcbi.1005595.

